# Graphene Micro-transistor Arrays Reveal Perfusion-Dependent Electrophysiological and Haemodynamic Signatures of Cortical Spreading Depolarizations in Ischaemic Stroke

**DOI:** 10.1101/2025.05.15.654197

**Authors:** S. Flaherty, A. Eladly, K. Hills, J. Merlini, E. Masvidal-Codina, E. Fernandez, X. Illa, E. Prats-Alfonso, E. Del Corro, J.A. Garrido, J. Meents, K. Kostarelos, S Allan, A. Guimerà-Brunet, R.C. Wykes

## Abstract

Cortical spreading depolarizations (CSDs) are large-scale disruptions of neuronal homeostasis that contribute to secondary injury in ischaemic stroke. In healthy cortex, CSDs induce vasodilation to meet metabolic demand, whereas in ischaemic tissue, they can provoke vasoconstriction and sustained hypoperfusion, exacerbating damage. We present a multimodal neurotechnology platform integrating flexible, transparent graphene solution-gated field-effect transistor (gSGFET) arrays with laser speckle contrast imaging (LSCI), enabling simultaneous DC-coupled electrophysiological recordings and cerebral blood flow imaging with high spatiotemporal fidelity. gSGFETs enable stable, distortion-free recording of infraslow potentials essential for resolving CSD waveforms in vivo. In two murine stroke models, we show that CSD waveform duration and morphology scale with local perfusion, delineating electrophysiological subtypes predictive of tissue viability and haemodynamic response. In metabolically compromised cortex, CSDs exhibit double peaks or negative ultraslow components that result in vasoconstriction, contrasting with narrow, monophasic waveforms eliciting vasodilation in healthy regions. Systemic ketamine shortens CSD duration and converts vasoconstriction to vasodilation in perfusion-deficit tissue, revealing a mechanism for its neuroprotective action. This platform enables high-fidelity investigation of brain–blood flow interactions in vivo.

## Introduction

Cortical spreading depolarizations (CSDs) are large, transient disruptions of neuronal and glial membrane potentials that propagate slowly through the cortex. First described by Leão in 1944 ^1^, CSDs are initiated in metabolically stressed but still viable brain tissue and are a hallmark of acute brain injury, particularly following focal cerebral ischaemia ^2,3^. These waves are characterised by near-complete breakdown of ionic homeostasis, resulting in neuronal and astrocytic depolarization, dendritic swelling, and spine distortion ^4,5^. Recovery from a CSD is energy-intensive, requiring ATP-dependent ion pumps to restore ionic gradients, a process that is feasible in healthy tissue but significantly impaired in ischaemic regions with reduced metabolic capacity ^3,6^.

CSDs have been implicated as a key mechanism driving lesion progression in ischaemic stroke. In the penumbra, the region surrounding the infarct core, recurrent spontaneous CSDs exacerbate tissue damage and correlate with infarct expansion ^7–9^. Rather than the expected hyperaemic response seen in healthy brain, CSDs in the ischaemic cortex can trigger prolonged vasoconstriction and hypoperfusion, a phenomenon termed “spreading ischaemia” ^10,11^. This inverse haemodynamic response impairs repolarisation and promotes a pathological feedback loop of repeated CSDs, calcium overload, and neuronal injury ^12,13^. Clustered CSDs pose a huge metabolic challenge for recovery and are thought to play a significant role in early cell death and lesion expansion.

Determining whether CSD waveform properties are correlated with localised tissue health and lesion progression in real time requires high-density, DC-coupled recordings free from attenuation and distortion. However, high-resolution electrographic mapping of CSDs remains technically challenging. Conventional electrode systems, even those capable of DC-coupled recordings, are limited by drift, impedance, and poor spatial fidelity ^14–16^. Graphene-based solution-gated field-effect transistors (gSGFETs) offer a promising alternative. These active transducers detect local voltage changes as current fluctuations across a graphene channel, enabling stable, wide-band recordings without the need for gate dielectrics. The intrinsic material properties of graphene; chemical inertness, flexibility, and high transconductance make gSGFETs well-suited for capturing the infraslow potentials associated with CSDs ^17–19^. These devices, with a total thickness of 10 µm, are compatible with simultaneous laser speckle contrast imaging (LSCI), allowing correlation of localised CSD waveforms with regional cerebral blood flow (rCBF) dynamics in real time.

In this study, we employ 16 and 30 channel epicortical gSGFET arrays for DC-coupled recordings in two well established preclinical models of acute focal cerebral ischaemia. We observe that CSD waveform duration and shape scale systematically with underlying perfusion gradients, delineating distinct electrophysiological subtypes that reflect local tissue viability. These electrophysiological signatures not only index tissue health but also predict the CSD-associated haemodynamic response, distinguishing CSDs that induce vasodilation from those that trigger vasoconstriction.

Ketamine has shown significant promise in modulating spreading depolarizations (SDs) in the context of ischaemic stroke and other forms of acute brain injury. The therapeutic administration of ketamine has emerged as a promising treatment to block or modulate CSDs in pre-clinical ^20–26^ and clinical ^27–30^ studies.

Using this integrated platform, we evaluated the capacity of ketamine to modulate CSD electrophysiology and associated haemodynamic responses during the early stages of stroke evolution. We demonstrate that systemic ketamine shortens CSD duration and converts inverse haemodynamic responses into vasodilation in metabolically compromised tissue, indicating a mechanism for the neuroprotective effects of ketamine in the post-stroke brain.

Together, these findings establish a technically advanced, multimodal platform for high-resolution, in vivo mapping of neurovascular dynamics, enabling unprecedented precision in characterizing CSD phenotypes and their vascular consequences in stroke. This provides a powerful and accessible tool for the neuroscience community to interrogate brain–blood flow interactions with high spatiotemporal fidelity.

## Material & Methods

### Animals

Mice were co-housed in ventilated cages, on a 12 h/12 h dark/light cycle, with food and water provided *ad libitum*. Animal procedures were conducted in accordance with the United Kingdom Animal (Scientific Procedures) Act 1986 and were approved by the Home Office. A total of 24 C57BL/6 mice were used in the described experiments. For all procedures, mice were weighed before being anaesthetized in an opaque induction chamber using 4% gaseous isoflurane (100% w/w; 988-3245, Henry Schein, USA) and medical gas (70% NO_2_ and 30% O_2_). Mice were then placed into a stereotaxic frame (David Kopf Instruments Ltd, USA). Lubrithal (Dechra, UK) was applied to each eye and pain relief, consisting of a subcutaneous Buprenorphine (0.03 mg/ kg, Ceva, France) injection and topical Emla cream (5% lidocaine/ prilocaine) (Aspen, UK) were administered. Iodine was applied to the surgical area. Anaesthesia was maintained using 1.5-2% isoflurane and medical gas was provided throughout the surgical and recording period. A rectal probe and heating pad was used to maintain an internal body temperature of 37ºc. During electrophysiological recording, the heating pad was turned off and a pre-heated (37ºc) Deltaphase isothermal pad (Braintree Scientific, USA) was used to maintain the body temperature. An incision was made down the midline and the skin retracted. The pericranium was removed and the surface of the skull cleaned with saline and dried.

### Craniotomy surgery

A 3 × 3 mm craniotomy was performed over the somatosensory cortex on the right hemisphere using a dental drill. To ensure minimal damage to the underlying tissue, ice-cold cortex buffered saline was continuously applied to the skull to prevent overheating during drilling. After the skull section was loose, the area was soaked in artificial cerebrospinal fluid (aCSF; mM concentrations in 500 ml distilled H_2_O: 125 NaCl, 4 KCl, 10 glucose, 10 HEPES, 2 CaCl_2_, 2 MgCl_2_, achieving a constant pH of 7.35) and the skull flap carefully removed using a fine needle. The underlying dura was left intact. To prevent the brain drying out, aCSF was applied after the procedure.

### Photothrombosis Induction of Focal Cerebral Ischaemia

Photothrombotic induction of focal cerebral ischaemia is a photochemical reaction that forms a reproducible thrombosis ^31^. In this work, photothrombosis was induced using a 561 nm laser (CL561-050, CrystaLaser, USA) with a power output of 50 mW before fibre and 35.5 mW at the fibre tip (PM16-130, Thor Labs, NJ, USA). An optic fibre cable (ThorLabs, M128L01, 400μm, USA) was positioned, using a micromanipulator, close to a corner of the gSGFET array. The tip of the optic fibre was enclosed with a custom-made black sheath housing to improve laser light localisation and reduce spread across the cortex. Prior to illumination, the photosensitive dye rose bengal (Sigma, UK), dissolved in saline solution (15 mg/ml), was slowly injected intraperitoneally. The animals were then kept in the dark to avoid non-specific rose bengal activation. The cortex was locally illuminated for an initial 60 second period. After full recovery of high frequency activity and the return of CBF to baseline, the cortex was illuminated, in the exact same area, for a further 180 seconds. The optic fibre was then removed from the preparation and electrographic and CBF monitoring continued for ~2 hours.

### Pharmacological CSD modulation

The therapeutic study consisted of two vehicle groups, group 1: IP Injection of saline (n = 15) and group 2: IP injection of ketamine (15 mg/kg) dissolved in saline solution (n = 7). Therapeutics were administered 5 seconds prior to the second 180 second illumination.

### Distal Middle Cerebral Artery Occlusion (dMCAo)

A straight vertical incision was made centrally between the ear and eye and the skin retracted to expose the underlying tissue/ muscle. The muscle underneath the suture line was superficially cut, detached from the skull, and retracted. The tissue covering the skull surface was removed and the area cleaned with saline. The skull was thinned above and around the vessel. Using a hooked needle tip, the thinned skull was carefully removed exposing the distal middle cerebral artery. The dura was left intact. The area was washed regularly with saline. Iron (III) Chloride (Sigma, UK) was dissolved in sterile water to form a 40% concentration solution (0.4g/ml). Filter paper was cut to a fine point and soaked in the ferric chloride solution for 10 seconds. The tip of the filter paper was placed directly onto the bifurcation of the vessel for 5 minutes. Occlusion was confirmed visually, by electrographic biomarkers and by a rapid perfusion decrease (~50% perfusion change) in the laser speckle contrast imaging.

### gSGFET Arrays

Flexible epidural neural probes (10 *μ*m in thickness) containing an array of 16 or 30 graphene micro-transistors were used. 16-channel (4 **×** 4 array, 400 *μ*m separation between transistors) and 30 channel (5×6 array, 500 *μ*m separation between transistors) arrays were used. gSGFET arrays were fabricated at the clean room facilities of IMB-CNM^17^. In brief, single-layer graphene was grown by chemical vapor deposition and transferred to a silicon wafer previously coated with a polyimide layer and patterned metal traces. After defining the graphene channels and before evaporating a second metal layer, UVO treatment was applied to improve the graphene-metal interface and reduce its contact resistance^32^. Finally, SU-8 was used as passivation layer and the polyimide layer was etched to define the geometry of the neural probes. Devices were gently peeled off from the wafer and inserted to zero insertion force connectors for electronic interfacing.

### Electrophysiological Recordings

Electrophysiological recordings were performed using flexible gSGFETs arrays. The tail of the array was inserted into a zero-insertion force (ZIF) connector on a printed circuit board (PCB) and lowered onto the dura using a micromanipulator. Signal acquisition and amplification was performed using one of two modified commercially available signal amplifiers: the Multichannel systems (MCS) ME2100 in vivo system (Multi Channel Systems MCS GmbH Germany) and the g.HIamp biosignal amplifier, (g.RAPHENE, g.tec medical engineering GmbH Austria). The MCS system required a custom made GE2100, gSGFET compatible, headstage (pre-amp). Wideband signals were acquired at 5000 Hz and 24-Bit. The g.tec system used a custom g.HIamp amplifier that acquired signals at 4600 kHz and 24 Bit. The g.HIamp system enabled simultaneous recording in two frequency bands with separate gains preventing amplifier saturation. Prior to recording, *I*_ds_ − *V*_gs_ curves were obtained at the start and end of experiments to determine the performance of the gSGFET array. A *V*_gs_ bias was selected at a voltage where transconductance was most optimal (steepest point of the *I*_ds_ − *V*_gs_ curve). Subsequent interpolation of acquired current signals into the transfer curve results in DC-coupled voltage signals. See supplementary figure 1Ai for further details^17^. A reference wire (Ag/AgCl_2_) was placed in the neck muscle.

**Figure 1.**
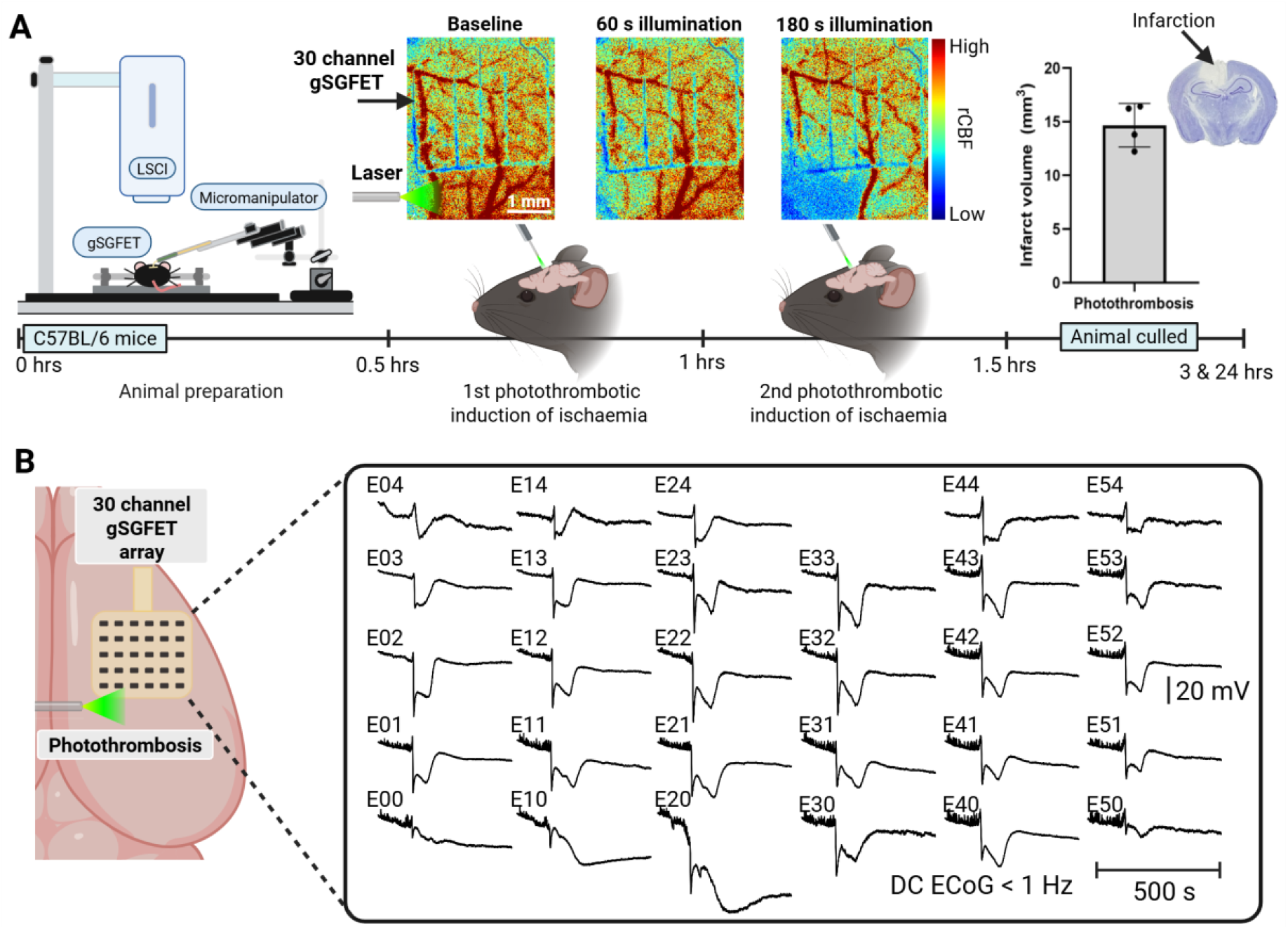
Experimental paradigm and recording configuration. **A**. Schematic of the experimental timeline for acute photothrombotic induction of focal cerebral ischaemia in anaesthetised mice. The setup enabled simultaneous DC-coupled electrophysiological recordings and laser speckle contrast imaging (LSCI) at the cortical surface beneath the transparent gSGFET array. Representative LSCI images display cortical blood flow (CBF) at baseline (pre-illumination), following a 60 second illumination (evoking a control CSD with minimal ischaemic insult) and after 180 second illumination (significant ischaemic insult). Animals were culled at timepoint 3 hrs after recording sessions. An additional cohort (n=4) were subjected to the same protocol without electrophysiological recordings and were culled after 24 hrs to assess infarct volume. **B**. Spatial map of DC-ECoG (<1 Hz) traces from 29 of 30 functional transistors arranged in a 5 × 6 grid configuration, demonstrating the spatial resolution of the 2.9 × 2.4 mm gSGFET array following the second (180 s) illumination.

### LSCI and Blood Oxygenation Monitoring

A Moor Instruments moorFLPI-2 Laser Speckle Contrast Imager (Moor Instruments, United Kingdom) was used for CBF monitoring. The stereotaxic frame and animal were carefully positioned onto the imaging base and the imager held above the preparation using a manipulator. The camera was focused onto the craniotomy and gSGFET array. The imaging field consisted of the right hemisphere, the midline and lambda. Images were acquired at 1 Hz (1 image per second) with a 2064 × 1544 resolution. To perform simultaneous CBF and blood oxygenation imaging, the above protocol was repeated however using the moorO2Flo (Moor Instruments, United Kingdom) imager. A read out of oxygenated haemoglobin (HbO) and deoxygenated haemoglobin (HbR) was acquired.

### Electrophysiology Data Analysis

Acquired electrophysiology recordings were analysed using Python 3.9 packages (Matplotlib 3.5.1, Numpy 1.21.5, Pandas 1.4.2, Seaborn 0.11.2, Neo 0.8.0) and the custom library PhyREC 0.5.1 and PhyREC 0.5.3. Transistor recordings were calibrated by interpolating the current signals into the corresponding branch of the *in vivo* measured transfer curve^17^. During recording calibration, down sampling was applied by a factor of 20. A full wideband signal is acquired in the experiment. In post-analysis, raw signals were filtered into 2 separate traces, the AC-ECoG component, and the DC-ECoG component. The AC-ECoG component was bandpass filtered 1-45 Hz. The DC-ECoG component was lowpass filtered < 1 Hz.

### CSD Event Detection

Semi-automated detection of CSD events was developed for this work, using Python scripts, to assist the analysis of large, multichannel, datasets. A bandpass filtered (0.05 – 0.001 Hz) signal derivative was applied to the DC-ECoG trace. At infraslow frequencies, only large voltage shifts (> 4 mV) in the DC-ECoG trace would cause large deflections in a 0.05 – 0.001 Hz bandpass filtered signal derivative (Fig. S3Ai-ii). This method of CSD event detection requires a stable signal. If a severe voltage drift is present in the recording and an offset correction applied, the CSD detection algorithm will detect the artefacts caused by offset correcting. All detected CSD events were manually checked before moving to the next step of the data analysis pipeline.

### CSD Parameter Extraction

The onset of an CSD was defined as the onset of a steep negative DC shift with a minimum amplitude threshold of −4 mV. For each CSD event, per transistor channel, the following parameters were extracted: Maximum CSD amplitude, CSD duration, and CSD depolarization speed. Maximum amplitude was defined by the peak voltage change from the baseline as an absolute voltage. CSD duration calculation commenced when the DC-ECoG voltage decreased below −4 mV and finished when the signal recovered above −4 mV. CSD depolarization speed was calculated by quantifying the slope of the DC shift. Python code available upon request.

### CSD Waveform Sorting

DC-ECoG CSD waveform sorting was manually conducted following strict criteria Single Peak (SP): A steep voltage decrease, below – 4 mV, that remains at or above a voltage maximum and then repolarises above – 4 mV, Double Peak (DP): A steep voltage decrease, below – 4 mV, at or above a voltage maximum, and then voltage decreases toward or beyond the voltage maximum. This is then followed by repolarisation above – 4 mV, Negative Ultraslow Potential (NUP): A steep voltage decrease, below – 4 mV, that remains at a voltage maximum and does not repolarise above – 4 mV and Hyper-inverted (HI): A positive deflection in channels that have previously recorded an NUP waveform type and when a SP or DP CSD is recorded in surrounding gSGFETs at the same time.

### Cerebral Blood Flow and Blood Oxygenation Analysis

16 or 30 (for 16 or 30 channel gSGFET arrays respectively) regions of interest (ROIs), measuring 500 *μ*m x 500 *μ*m, were defined over each transistor in post analysis using the moorFLPI-2 Review software (V5.0). All perfusion values were then exported as .Txt files and loaded into Python 3.9 packages (Matplotlib 3.5.1, Numpy 1.21.5, Pandas 1.4.2, Seaborn 0.11.2, Neo 0.8.0). Baseline cerebral blood flow (CBF) was monitored for approximately 10 minutes before induction of focal cerebral ischaemia. In order to normalise rCBF data, a stable reference point was selected during the 10-minute baseline recording. All rCBF data in this work is presented as normalised perfusion values, with the units ‘perfusion units’ (PU). CSD induced haemodynamic responses were manually sorted and defined by: vasoconstriction (VC), a normalised perfusion decrease below baseline and increased toward baseline during or after a CSD; biphasic (BP), a normalised perfusion decrease below baseline and increased beyond baseline during or after a CSD; vasodilation (VD), normalised perfusion increase above baseline and decreased towards baseline during or after a CSD. During illumination time windows, when inducing photothrombosis, an artefact was present in the LSCI, causing the rCBF data to be skewed and unusable. Manual laser blanking was applied to remove this data. Blood oxygenation HbO and HbR data was exported in the same manner as described for rCBF data. HbO and HbR recovery time was manually calculated.

### Analysis and Statistics

GraphPad Prism 10.4 software was used to perform statistical analysis. Statistical tests are described in the figure legends or results text (mean ± sd). For pairwise comparisons, a two-sided unpaired t-test was used. Data sets greater than two groups were compared using either one-way ANOVA or two-way ANOVA. A Spearman’s rank correlation coefficient or simple linear regression was used to describe the correlation between two variables. The number of animals and p-values are described in the figure legend or results text.

## Results

### Integrated multimodal platform enables simultaneous DC-coupled electrophysiology and cerebral blood flow imaging

To investigate the relationship between CSDs, CBF, and tissue viability, we developed a multimodal platform combining DC-coupled graphene solution-gated field-effect transistor (gSGFET) arrays with LSCI (Fig. 1A). This setup enabled high-resolution mapping of both neuronal activity and rCBF in anesthetised mice undergoing focal cerebral ischaemia. The transparent, flexible gSGFET array was placed epidurally over the cortex, allowing for simultaneous electrophysiological recording and full-field optical imaging (Fig. S1Ai-ii). Prior to recordings, transistor-specific transfer curves were obtained to determine the optimal gate bias (Vgs) for maximizing current sensitivity (Ids)^17^. The DC-stable acquisition system preserved infraslow potentials crucial for resolving CSD waveforms (Fig. 1B). LSCI, focused through the transparent array, provided concurrent surface rCBF measurements with high spatiotemporal resolution (Fig. S1Bii).

### CSD duration is inversely correlated with regional perfusion following ischaemic injury

We first induced focal cerebral ischaemia using the photothrombotic (PT) model (Fig. 1A), with validation in the dMCAo model (Fig. S2A). After an initial brief photothrombosis (60s illumination), a single CSD was reliably evoked, but duration did not vary with perfusion level (Fig. S4Ai-ii). The multichannel array enables precise quantification of spatial variations in CSD properties across the cortex. (Fig. 2A). Note that after the second illumination (additional 180s) the duration and amplitude of SDs diminish with increasing distance from the site of ischaemic induction, and that clusters of SDs can also be observed in this region. This extended illumination, results in substantial ischaemia developing adjacent to the recording array, producing a gradient of perfusion across the cortex (Fig. 2B). Under these conditions, CSDs varied significantly in duration across the array and were strongly dependent on underlying rCBF (Fig. 2Bii-C). CSD duration negatively correlated with normalized perfusion (Spearman’s r = −0.65, p = 2.1e^−25^, n = 15) (Fig. 2Di). Stratifying CSDs by local perfusion revealed progressively shorter durations with increasing rCBF: 144.8±26.8 s at 0.5 PU; 104.7±39.4 s at 0.7 PU; 66.2±33.9 s at 0.9 PU; and 51.5±21.9 at 1.1 PU (One-way ANOVA, **** *p* < 0.0001) (Fig. 2Dii).

**Figure 2.**
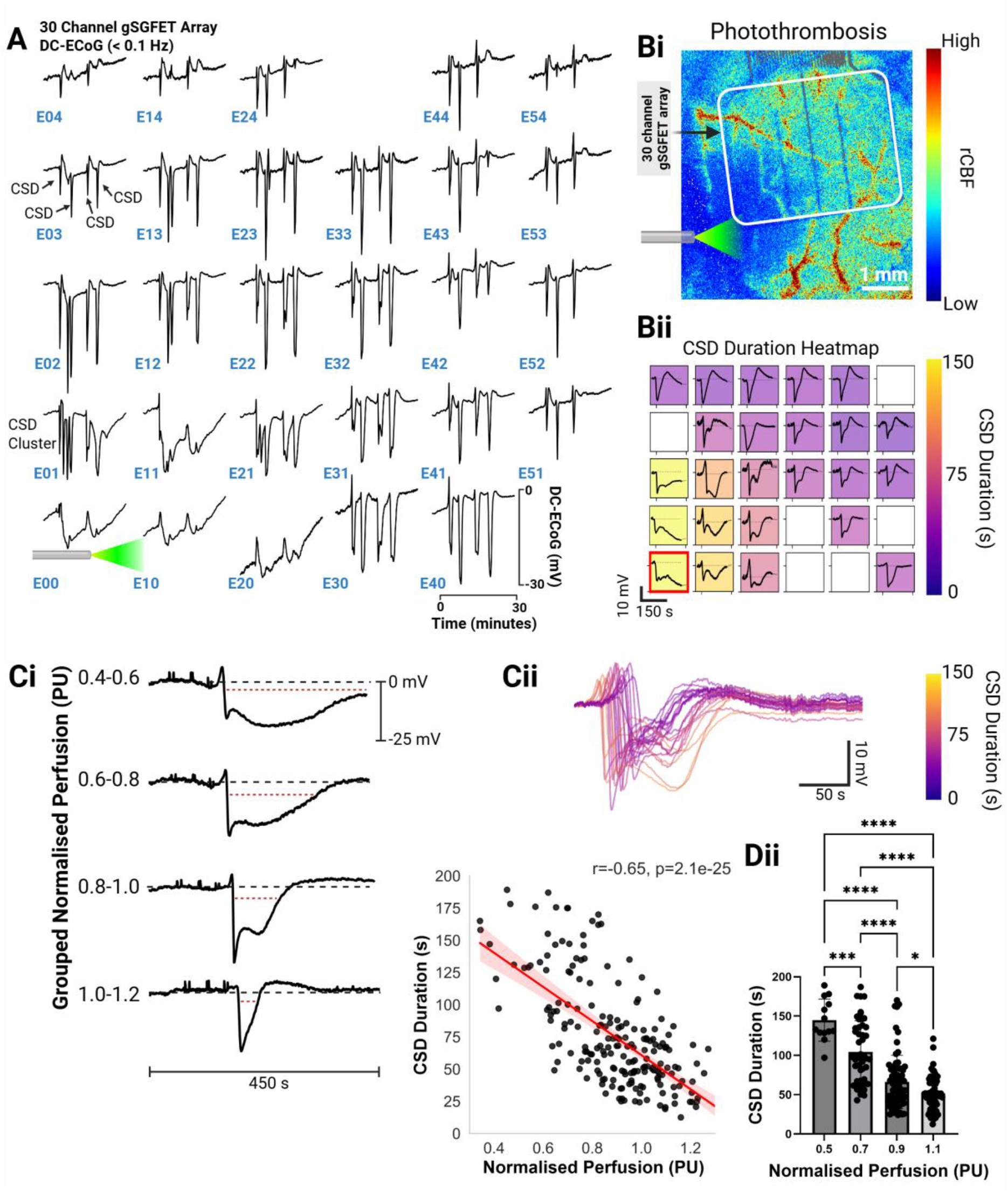
Duration of ischaemia-induced cortical spreading depolarizations (CSDs) is dependent on regional cerebral blood flow (rCBF). **A**. An example 30 min DC-ECoG (< 0.1 Hz) recording acquired using a gSGFET array following 180 s photothrombotic induction of focal cerebral ischaemia (green laser) in anaesthetised mice. 28 of 30 active transistors illustrate the spatial and temporal mapping of CSD propagation across the cortical surface **Bi**. Single frame laser speckle contrast imaging (LSCI) showing resulting perfusion deficit (dark blue region) following 180 s photothrombosis. The recording gSGFET is highlighted by a white box. **Bii**. Heatmap of CSD duration (s) for a single CSD event recorded in the same preparation as Bi. Empty boxes indicate non-functioning channels; the red outline marks the channel nearest the lesion core. **Ci**. Representative CSD traces recorded in regions with differing normalised perfusions (PU), illustrating perfusion-dependant variability in CSD duration. The dotted redline indicates the change in CSD duration. **Cii**. Overlay of 29 CSD traces from a single CSD event recorded across a 30 channel gSGFET array. Trace colour represents CSD duration (s). **Di**. Scatter plot displaying a significant negative correlation between CSD duration (s) and underlying normalised perfusion (PU) (Spearman’s r = −0.65, p=2.1e^−25^). **Dii**. Grouped analysis of mean CSD duration (s) as a function of normalised perfusion (PU), revealing significant differences across most perfusion bins (mean ± s.d.).

This relationship was corroborated in the dMCAo model, where topical ferric chloride produced a rapid occlusion and CSD onset (Fig. S5A). A similar inverse correlation between CSD duration and perfusion was observed (Spearman’s r = −0.68, p = 1.1e-32, n = 15) (Fig. S5Bi-ii), confirming that CSD duration reflects the underlying metabolic state across models.

### CSD waveform morphology is shaped by local metabolic state

The spatial resolution of the gSGFET array allowed classification of CSD waveform morphologies within a single preparation (Fig. 3A). Across both stroke models, four distinct waveform types were identified: single peak (SP), double peak (DP), negative ultra-slow potential (NUP), and hyper-inverted (HI) (Fig. 3B). HI is a positive deflection observed in channels that previously recorded an NUP waveform, coinciding with the detection of DP or SP-type CSDs in adjacent gSGFETs. HI were only observed in depolarized tissue. The waveform types showed an association with local perfusion. In the PT model, SP CSDs were recorded at 0.96 ± 0.18 PU, DP at 0.70 ± 0.23 PU, NUP at 0.42 ± 0.18 PU, and HI at 0.48 ± 0.15 PU (One-way ANOVA, **** *p* < 0.0001) (Fig. 3Ci-iii). DP waveforms were significantly longer in duration than SP (130.0 ± 59.0 s vs. 50.89 ± 21.92 s, **** *p* < 0.0001) (Fig. 3Cii-Civ).

**Figure 3.**
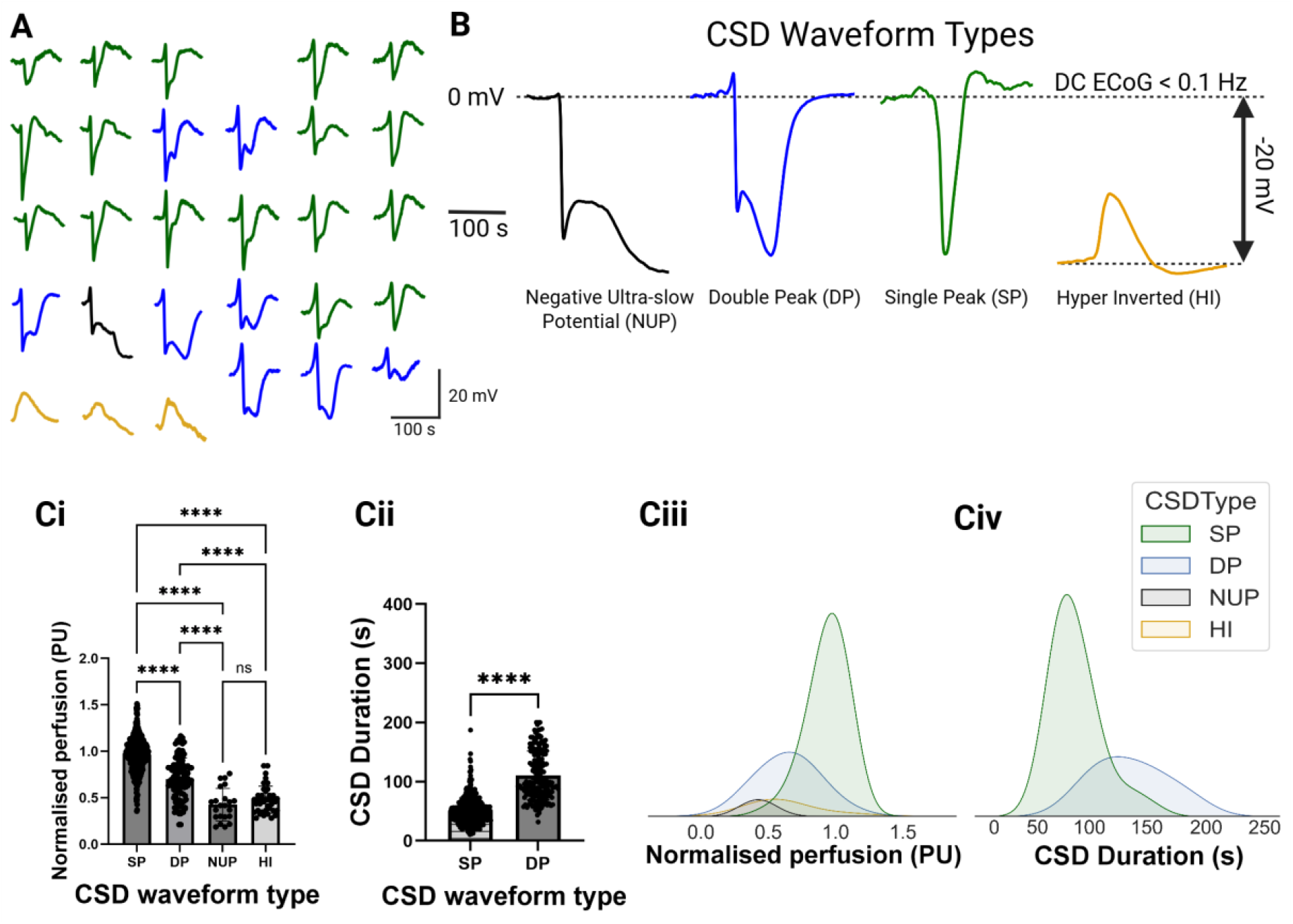
Cortical spreading depolarization waveforms are dependent on regional cerebral blood flow (rCBF). **A**. Representative CSD waveform types recorded across 29 out of 30 functional gSGFET channels in response to photothrombotic induction of focal cerebral ischaemia. **B**. Four distinct CSD waveform types were identified: Negative ultra-slow potential (NUP), double peak (DP), single peak (SP) and hyper inverted (HI). **Ci**. Plot displaying CSD waveform types (457 SP, 113 DP, 22 NUP and 39 HI waveforms) as a function of mean normalised perfusion (PU) demonstrating perfusion-dependant waveform types **Cii**. Comparison of mean SP and DP CSD (535 SP and 192 DP CSDs) duration (s) highlighting prolonged durations associated with DP CSDs. **Ciii**. Density plots showing the distribution of CSD waveform types across a range of normalised perfusions (PU). **Civ**. Density plot of SP and DP CSD duration (s) distribution.

In the dMCAo model, similar trends were observed: SP at 0.80 ± 0.16 PU, DP at 0.65 ± 0.16 PU, NUP at 0.59 ± 0.12 PU, and HI at 0.51 ± 0.08 PU (Fig. S6Ai-iii). DP waveforms again had significantly longer durations than SP (129.90 ± 58.97 s vs. 53.32 ± 20.47 s, unpaired t-test, **** *p* < 0.0001) (Fig. S6Aii-iv).

### CSD-induced haemodynamic responses reflect perfusion status

To understand how CSDs influence local haemodynamics, we classified CSD-evoked rCBF changes into vasoconstrictive (VC), biphasic (BP), or vasodilatory (VD) responses (Fig. 4A). In the PT model, response types occurred at significantly different perfusion levels: VC at 0.79 ± 0.20 PU, BP at 0.85 ± 0.18 PU, and VD at 1.01 ± 0.18 PU (One-way ANOVA, **** *p* < 0.0001) (Fig. 4Bi-ii). CSD-induced vasodilatory responses were more frequently associated with SP waveforms, whereas DP, HI and NUP CSDs were predominantly linked to VC and BP haemodynamic responses (Fig. 4Biii).

**Figure 4.**
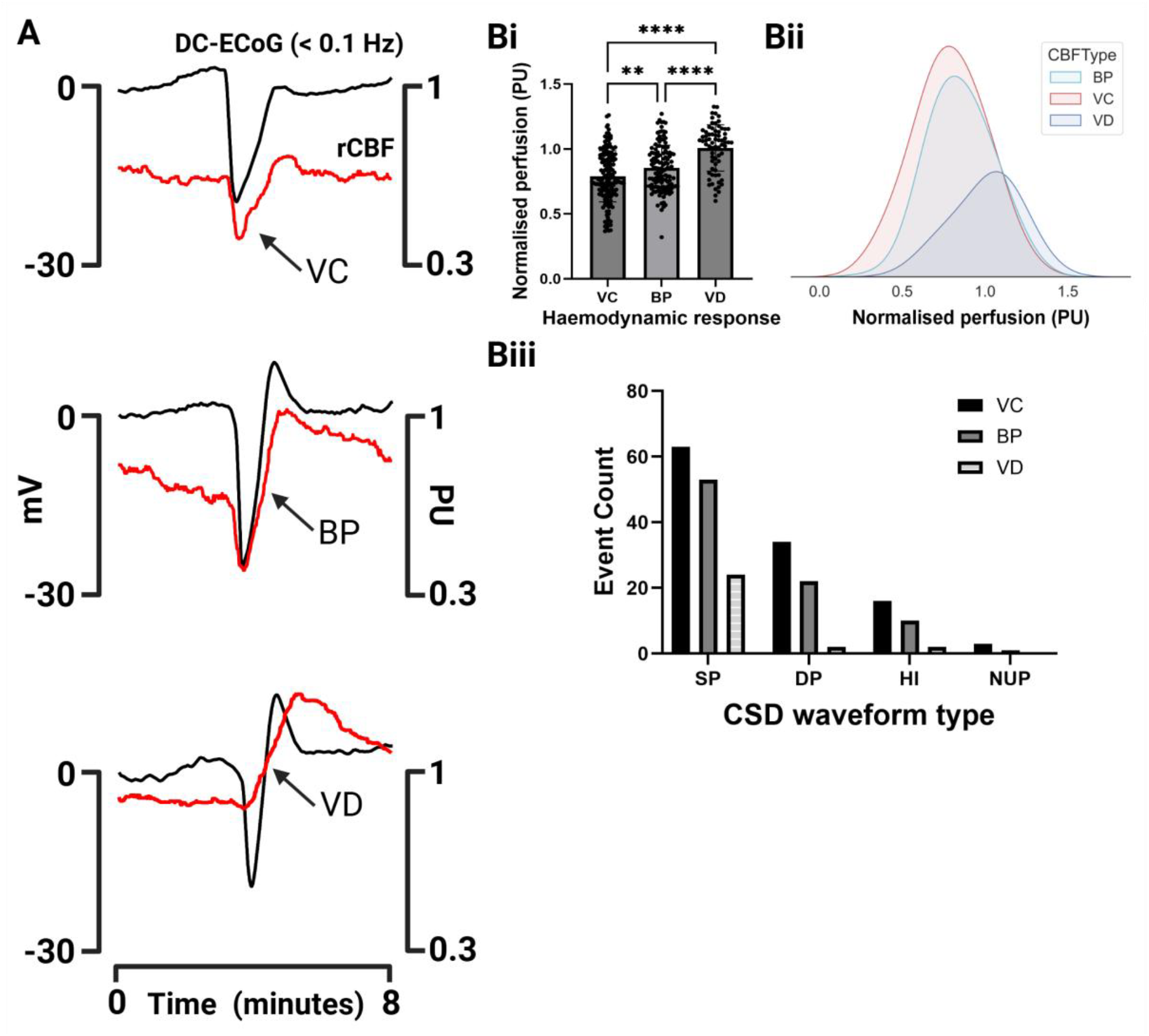
Cortical spreading depolarization induced haemodynamic responses are dependent on regional cerebral blood flow. **A**. Example DC-ECoG and rCBF traces post photothrombosis in anaesthetised mice with labelled CSD induced haemodynamic response. Three haemodynamic responses were observed in this work, vasoconstriction (VC), biphasic response (BP) and vasodilation **Bi-ii**. CSD-induced haemodynamic response type plotted in function of mean normalised perfusion (PU) and presented as a density distribution plot. **Biii**. Count of CSD-induced haemodynamic responses with respect to individual CSD waveform types (SP induced 63 VC, 53 BP and 24 VD; DP induced 34 VC, 22 BP and 2 VC; HI induced 16 VC, 10 BP and 2 HI and NUP induced 3 VC and 1 BP).

Comparable patterns were observed in the dMCAo model (Fig. S8A). CSD-evoked VC responses were recorded at 0.59 ± 0.17 PU, BP at 0.75 ± 0.18 PU, and VD at 0.74 ± 0.08 PU (One-way ANOVA, **** *p* < 0.0001) (Fig. S8B). These findings indicate that CSD-associated hemodynamic responses shift from vasodilation to vasoconstriction with increasing metabolic compromise.

### Local Haemodynamic and Oxygenation Changes Associated with CSDs

Given the high metabolic demand of CSDs, we monitored localized haemoglobin oxygenation levels to assess their contribution to disruptions in tissue oxygenation. We applied our multimodal recording setup to simultaneously monitor DC-ECoG, rCBF, and haemoglobin oxygenation dynamics (HbO and HbR; Fig. S9A). We quantified the recovery time of HbO and HbR following CSD onset (Fig. S9B). HbO recovery time exhibited a strong inverse correlation with normalized perfusion (PU), indicating that impaired perfusion delays reoxygenation. In contrast, HbR recovery times showed a weaker association with perfusion.

CSD duration was significantly correlated with the recovery time of both HbO (*r*^*2*^ = 0.59, *p* < 0.0001) and HbR (*r*^*2*^ = 0.25, *p* < 0.0001; Fig. S9D), indicating that prolonged depolarizations impose a sustained metabolic burden. Distinct CSD waveform morphologies were also predictive of oxygen dynamics. Specifically, NUP and DP waveforms were associated with extended HbO depletion (Fig. S9Ei). While HbR recovery was less tightly coupled to waveform features, NUP events exhibited significantly prolonged HbR recovery times relative to other waveform classes (Fig. S9Eii). These findings highlight the utility of CSD waveform profiling in forecasting the spatiotemporal evolution of metabolic stress in vulnerable tissue. SD-induced changes in blood oxygenation have been observed clinically, with a study reporting that the absence of a change in blood oxygenation response to CSD correlated with poor outcomes in patients with malignant stroke^33^. While we did not observe this association, this probably reflects the earlier timepoint of our measurements and the comparatively less severe pathology in our rodent model.

### Ketamine shortens CSD duration and modulates waveform

To explore the effects of modulating CSD properties, we investigated the impact of ketamine (15 mg/kg IP) (Fig. 5A). We used a dose that is reported as non-sedative in awake mice (Masvidal et al 2021). Following ketamine administration, CSD waveforms were shorter when compared to saline treated animals (Fig. 5B). Comparison of CSD durations recorded after the initial 60 s photothrombosis displayed no significant difference between groups (Fig. 5Ci). However, after injecting ketamine and illuminating for 180 s, CSD duration no longer correlated significantly with perfusion (Spearman’s r = −0.14, p = 0.02, n = 7), in contrast to the robust negative correlation observed in saline-treated animals (Spearman’s r = −0.65, p = 2.1e^−25^, n = 15) (Fig. 5Cii). CSD durations in ketamine-treated animals were significantly shorter at each perfusion level, notably durations reduced to 49.6 ± 43.7 s at 0.5 PU compared to 144.8 ± 26.8 s in saline controls and 39.6 ± 24.2 at 0.7 PU compared to 104.7 ± 39.4 (two-way ANOVA, **** *p* < 0.0001). SP waveform types were more common following ketamine administration displaying a reduction in longer DP CSDs (Fig. 5Di-ii). The number of spontaneous CSDs did not change between groups following 180 s illumination (Fig. 5Ei). CSD propagation speed in ketamine animals was comparable to saline-controls for the first and second CSDs but then began to significantly differ with subsequent CSDs (Fig. 5Eii).

**Figure 5.**
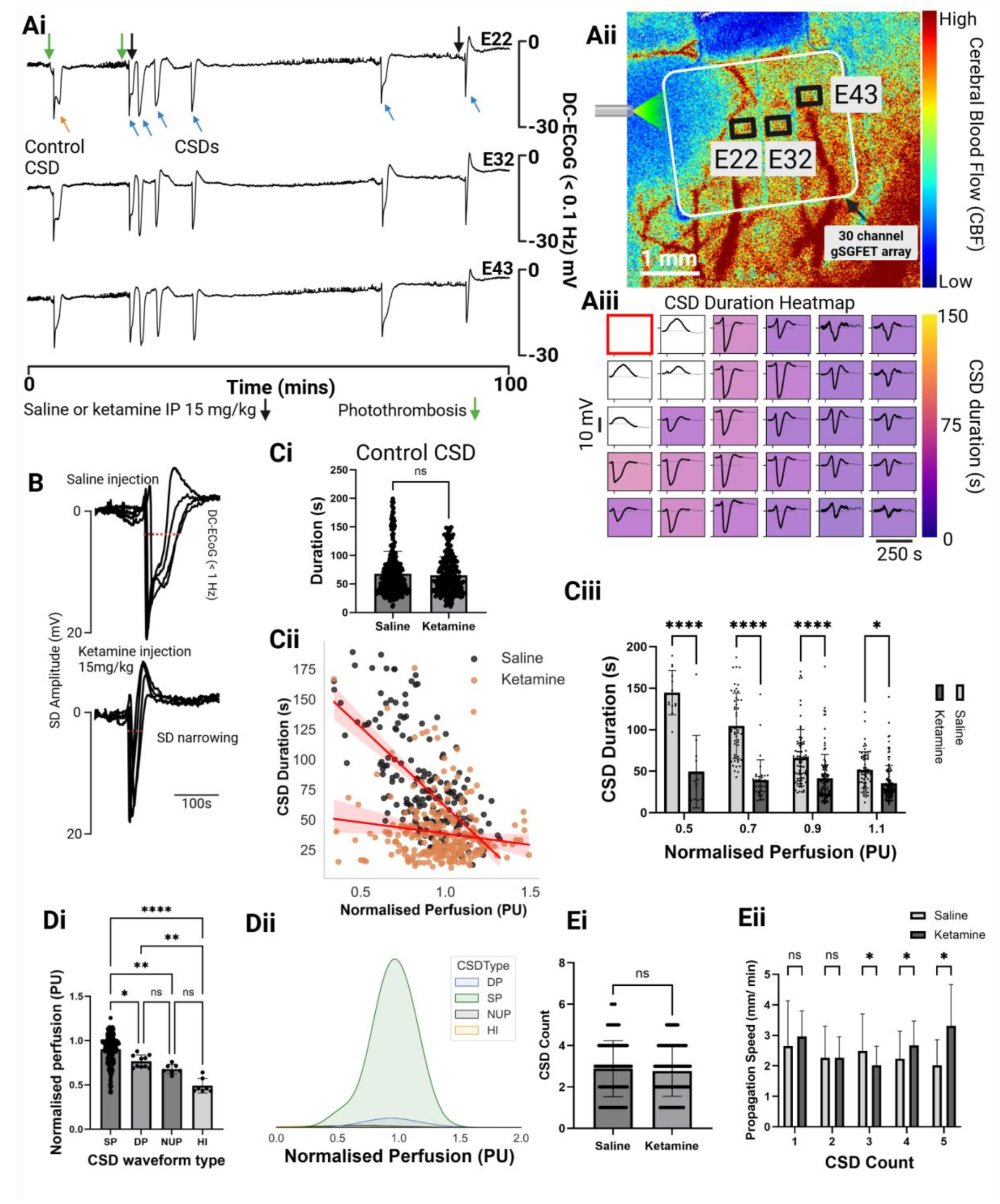
Ketamine modulation of cortical spreading depolarization (CSD) duration and waveform. **A**. Three channel 100 mins DC-ECoG recording. 1^st^ 60 seconds and 2^nd^ 180 seconds photothrombosis marked by green arrows. Saline or 15mg/kg ketamine was administered via IP injection moments after the 2^nd^ illumination (black arrow) **Aii**. Single frame from Laser speckle contrast imaging (LSCI) characterising an area of cerebral ischaemia following photothrombosis (dark blue region) and 3 highlighted gSGFETs at 0.5 mm separation. The selected channel recordings from this experiment are plotted in Ai. **Aiii**. CSD duration (s) displayed as a 5×6 heatmap. Each trace is 250 s in length. B. Four overlaid CSD traces from a saline injected animal and a ketamine injected animal. Note the reduction in CSD duration (s). **Ci**. Mean control CSD duration (s) following 1^st^ 60 s illumination from saline and ketamine groups. **Cii**. Comparison of the correlation between CSD duration (s) and normalised perfusion (PU) between saline injected animals (r = −0.65, p=2.1e^−25^) and ketamine injected animals (r= −0.14, p= 0.02) post photothrombosis. **Ciii**. Plot of CSD duration (s), from saline and ketamine injected animals, in function of normalised perfusion (PU). **Di-Dii**. CSD waveform types extracted from ketamine injected animals plotted against mean normalised perfusion (PU), shown as a bar and kernel density estimate plot. **Ei**. Number of CSDs recorded post injection of saline or ketamine within the first 80 minutes following 180 s photothrombosis. **Eii**. Comparison of saline and ketamine CSD propagation speed in function of CSD count following 2^nd^ 180 s photothrombosis.

### Ketamine prevents CSD-induced inverse haemodynamic responses

In saline-treated animals, CSDs in metabolically compromised regions predominantly evoked vasoconstriction (Fig. 6A). Administration of ketamine markedly attenuated these vasoconstrictive responses (Fig. 6Bi-ii). Of the 366 responses recorded in ketamine treated animals, 343 were VD and 23 BP. In contrast, saline treated animals exhibited a broader distribution (432 responses), of which 71 were VD, 154 BP and 207 VC. Representative traces confirmed the conversion of vasoconstrictive to vasodilatory responses even in low-perfusion regions, suggesting that ketamine restores neurovascular coupling and suppresses the pathological inverse haemodynamic response to CSD. Resting rCBF, assessed 1.5 hours after the 180 s illumination, was significantly elevated in ketamine-treated animals relative to the distance from the lesion core (Fig. 6C). While normalised perfusion declined following multiple CSDs in saline-treated animals (Fig. 6Dii), ketamine treatment led to a sustained increase in perfusion across successive CSDs (Fig. 6Dii). Notably, groupwise analysis of the 3^rd^ CSD revealed significantly enhanced perfusion at 1.5 and 2.0 mm from the lesion in the ketamine cohort (Fig. 6Diii). Collectively, these findings underscore the capacity of our gSGFET-LSCI platform to resolve dynamic neurovascular responses with high spatiotemporal fidelity and identify ketamine as a potent modulator of CSD-induced haemodynamics in the ischaemic brain.

**Figure 6.**
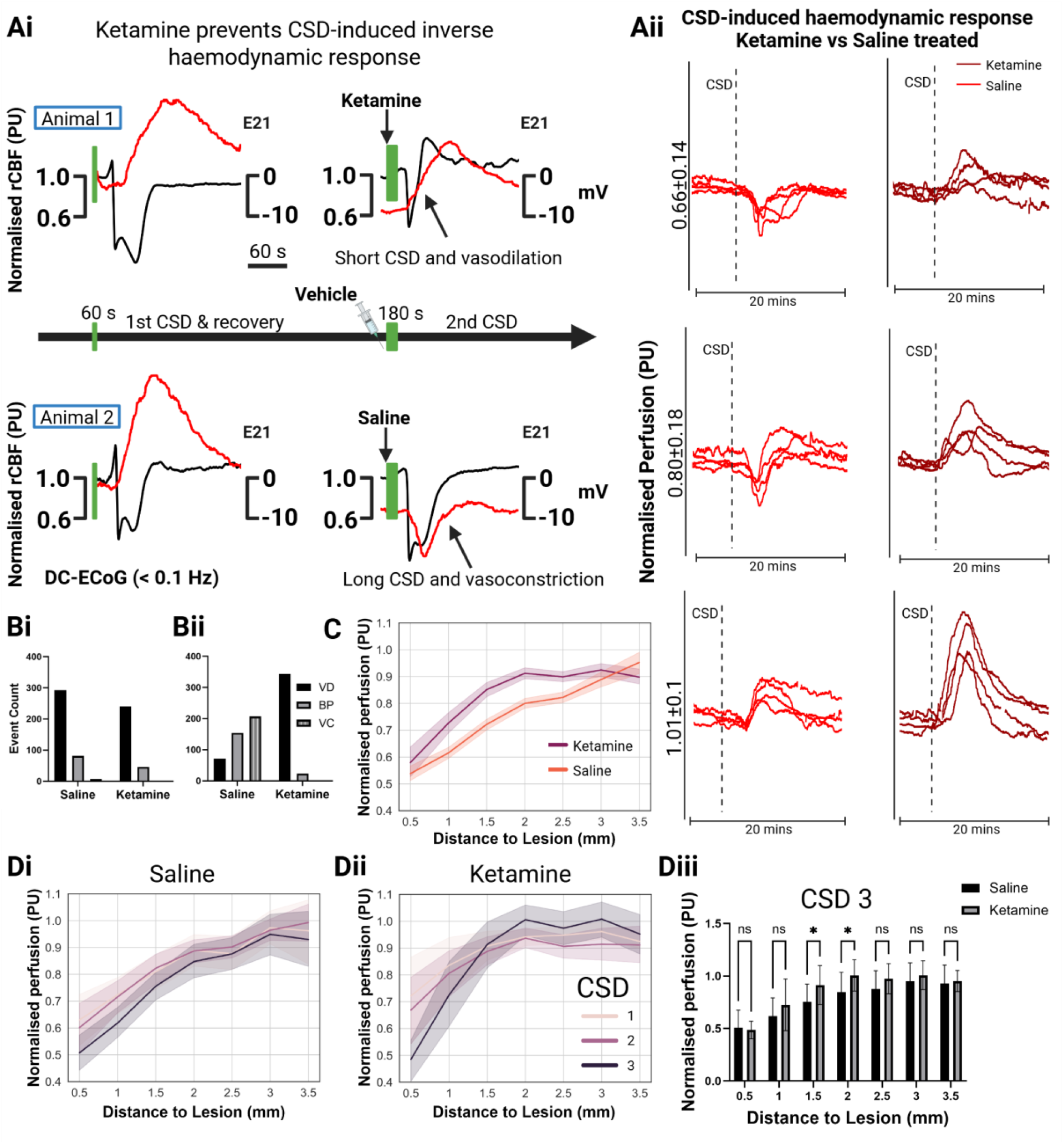
Ketamine prevents the inverse haemodynamic response to spreading depolarizations. **Ai** DC-ECoG and rCBF (PU) traces from saline and ketamine (15 mg/kg) injected animals after 1^st^ 60 s photothrombosis (control) and after 2^nd^ 180 s photothrombosis. Green bars illustrate illumination time and black arrow marks time of vehicle administration. A**ii**. rCBF traces from saline and ketamine injected animals displaying example CSD-induced haemodynamic responses at similar normalised perfusion ranges (PU). Black dashed lines mark the start of the CSD. **Bi-ii**. CSD-induced haemodynamic response type from saline and ketamine injected animals following 1^st^ 60 s and 2^nd^ 180 s photothrombosis respectively. **C**. Final resting (minimal influence from CSD induced perfusion changes) rCBF (PU) in function of distance to lesion core (mm) quantified 1.5 hours following 2^nd^ 180 s photothrombosis from saline and ketamine injected animals. **Di-ii**. Normalised perfusion (PU) quantified at distance (mm) from the lesion core with respect to CSD number following 2^nd^ 180 s photothrombosis from saline and ketamine injected animals respectively. **Diii**. Quantification of normalised perfusion (PU) against distance (mm) to lesion core. Plot compares CSD 3 from saline and ketamine injected animals.

## Discussion

Our study establishes a multimodal platform that integrates flexible graphene-based micro-transistor arrays with LSCI to resolve CSDs and their associated vascular responses with high spatiotemporal fidelity. By enabling stable, DC-coupled electrophysiological recordings in parallel with real-time CBF imaging, this approach overcomes long-standing technical limitations in studying CSD dynamics and their contribution to secondary injury in stroke.

We show that CSD waveform features, specifically duration and waveform scale with regional perfusion, reflecting local tissue viability. In metabolically intact cortex, CSDs are coupled to vasodilation. In contrast, ischaemic regions exhibit prolonged, complex waveforms including biphasic peaks and ultraslow components, frequently accompanied by vasoconstriction. These distinct electrophysiological phenotypes index the tissue’s metabolic capacity and predict the ensuing haemodynamic response, supporting the hypothesis that CSD waveform shape serves as a real-time biomarker of regional tissue health. A continuum of CSD duration associated with haemodynamic responses, ranging from vasoconstriction in ischaemic tissue to vasodilation in healthy tissue, has been recognized^10^. The key advantage of our approach and technology is the ability to capture this full spectrum with high spatial and temporal resolution using a single recording gSGFET array.

We demonstrate that systemic administration of ketamine shortens CSD duration and converts vasoconstrictive responses into vasodilation in perfusion-deficient tissue. This modulatory effect reveals a mechanistic basis for ketamine’s neuroprotective properties in acute brain injury. By interrupting the pathological feedback loop of spreading ischaemia, energy failure, and recurrent CSDs, ketamine may preserve tissue viability in the ischaemic penumbra. These findings align with prior preclinical and clinical reports of ketamine’s efficacy in suppressing SDs and mitigating lesion expansion ^20–23,27–30,34^ while offering new insights into its effects on CSD haemodynamic coupling.

Future studies should explore how CSD waveform phenotypes evolve over time, particularly in the subacute phases of stroke. Additionally, integrating this platform with calcium imaging^35^, optogenetics ^18,36–38^ or metabolic biosensors could deepen our understanding of the cellular mechanisms that govern CSD initiation, propagation, and resolution, as well as their regional impact on tissue health under pathological conditions.

Future clinical translation of this technology could enable real-time, high-fidelity monitoring of lesion evolution in patients with acute brain injury, providing a functional biomarker of tissue viability. By improving the detection and characterization of spreading depolarizations, this approach may support dynamic assessment of injury progression and allow for the stratification of patient populations most likely to benefit from targeted neuroprotective interventions ^19,39^,

In summary, our findings define distinct electrophysiological signatures of CSDs that encode tissue state and vascular response in ischaemic stroke. By leveraging high-resolution, multimodal recordings, we identify ketamine as a potent modulator of CSD waveform dynamics and cerebrovascular outcome, advancing the framework for therapeutic intervention in acute brain injury.

## Data availability

All relevant data obtained to evaluate the main findings of the paper are openly available from the corresponding author upon reasonable request.

## Acknowledgements

This research was funded by the European Union’s Horizon 2020 research and innovation programme under grant agreement no. 881603 (Graphene Flagship Core 3) and the European Union’s Horizon. This work was supported by the Medical Research Council [Grant Ref: MR/Y014545/1].

