## Supplementary Material for "Graphene Micro-transistor Arrays Reveal Perfusion-Dependent Electrophysiological and Haemodynamic Signatures of Cortical Spreading Depolarizations in Ischaemic Stroke"

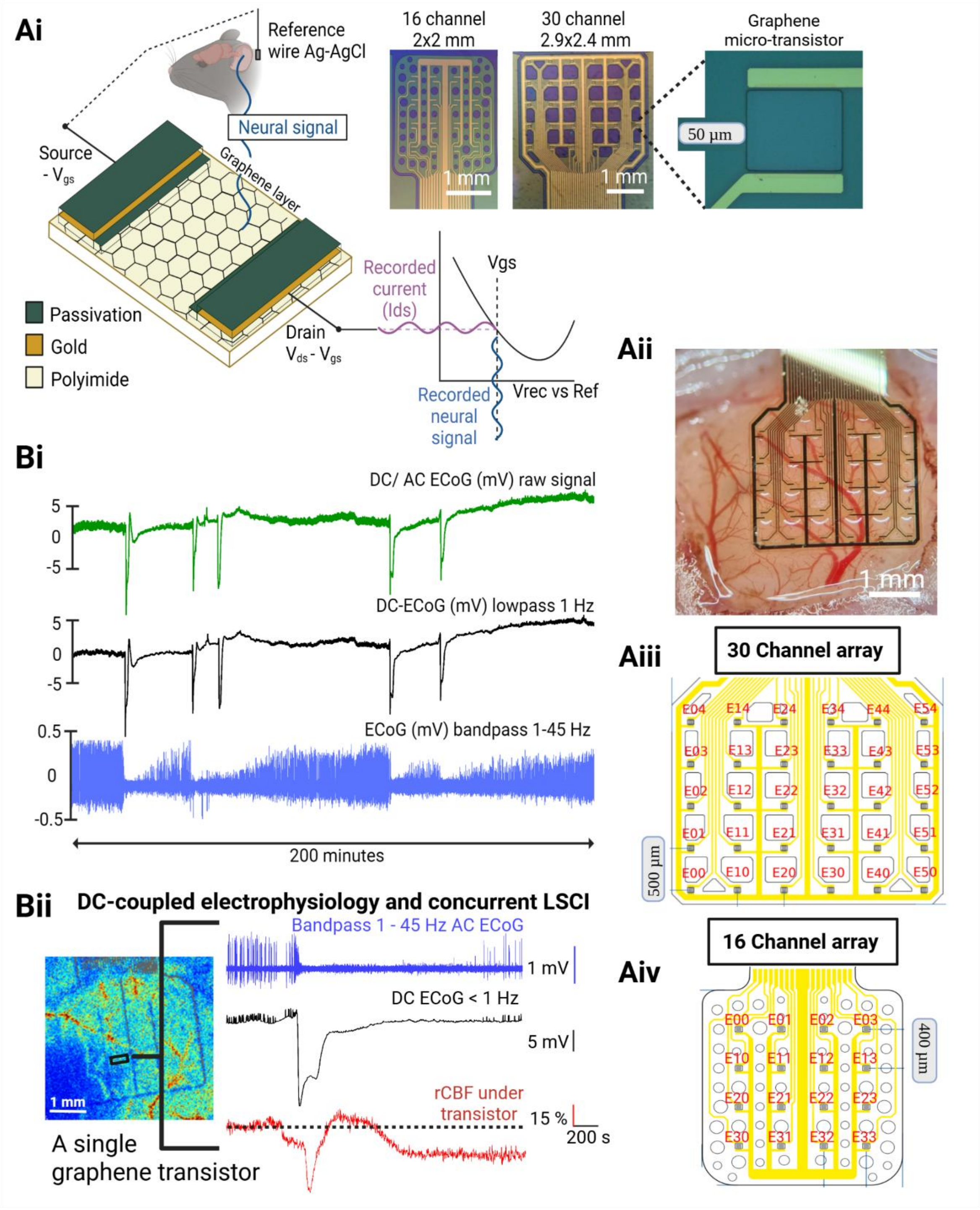


Supplementary figure 1. **Ai**. Schematic of a single graphene solution-gated field-effect transistor (gSGFET) and optimum voltage gate source (Vgs) biasing. Optical images of a 16 and 30 channel gSGFET array, including a blow up of a single gSGFET. **Aii**. Optical image of a 30 channel gSGFET array on the surface of a mouse cortex (dura intact). **Aiii-iv**. Schematic of gSGFET channel map for a 30 channel and 16 channel array respectively. **Bi**. Example 200-minute wide-bandwidth recording with a gSGFET array. Green trace is the raw ECoG signal, black trace the lowpass (1 Hz) DC-ECoG and blue is the bandpass (1-45 Hz) AC-ECoG. **Bii.** AC-ECoG (blue trace 1-45 Hz) and DC-ECoG (black trace < 1 Hz) electrophysiology with concurrent laser speckle contrast imaging rCBF monitoring from a single gSGFET.

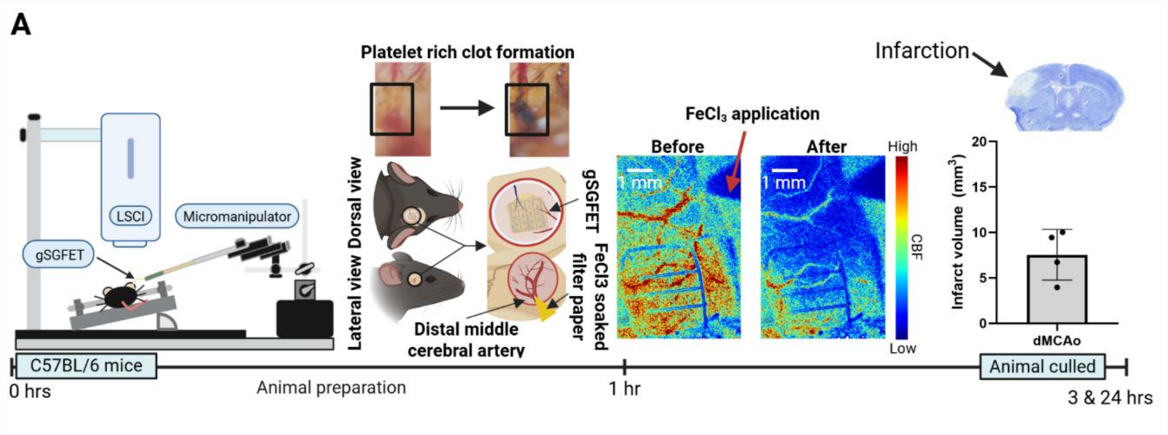


Supplementary figure 2. **A**. Schematic of the experimental timeline for acute dMCAo induction of focal cerebral ischaemia in anaesthetised mice. The setup enabled simultaneous DC-coupled electrophysiological recordings and laser speckle contrast imaging (LSCI) at the cortical surface beneath the transparent gSGFET array. Representative LSCI images display cortical blood flow (CBF) at baseline (before dMCAo), and after dMCAo. Animals were culled at timepoint 3 hrs after recording sessions. An additional cohort (n=4) were subjected to the same protocol without electrophysiological recordings and were culled after 24 hrs to assess infarct volume.

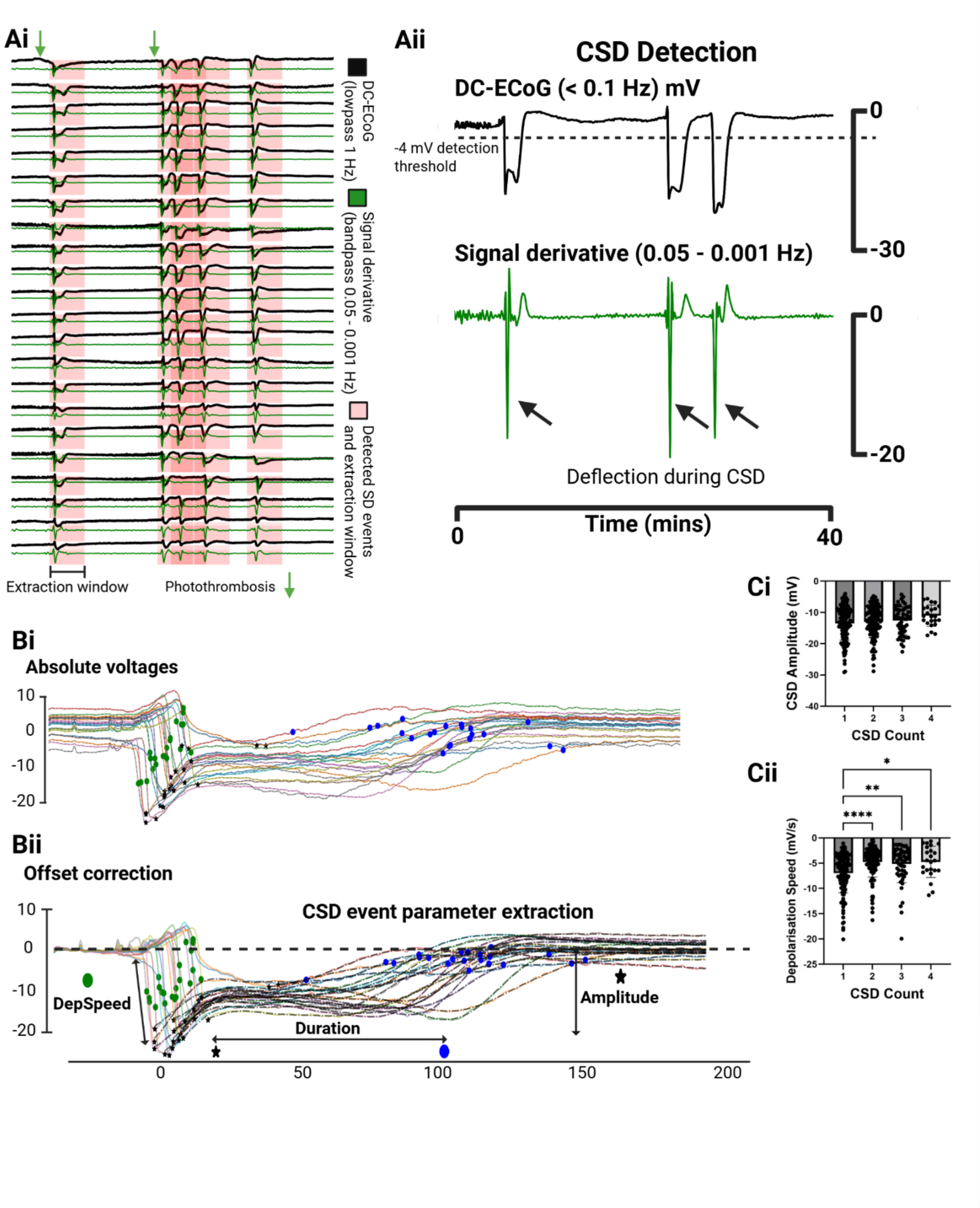


Supplementary figure 3. **Ai**. Semi-automated detection of DC shifts. Black trace is DC-ECoG (lowpass 1 Hz), and green trace is the signal derivative (bandpass 0.05 – 0.001 Hz). Pink boxes highlight detected SD and extraction window (-30 to 250 s). **Aii**. Blow up of a single DC-ECoG trace and signal derivative. **Bi-ii**. Absolute voltage extraction of a single spreading depolarisation event and the same event displayed underneath with an offset correction to 0 mV. Extracted parameters: Depolarisation speed (green circle), amplitude (black star), end of the event (blue circle) and duration (time between black star and blue circle). **Ci-ii**. Quantified CSD amplitude (mV) and depolarisation speed (mV/s) plotted in function of CSD count following 180 s illumination (One-way ANOVA, **** $p < 0.0001$, n =15).

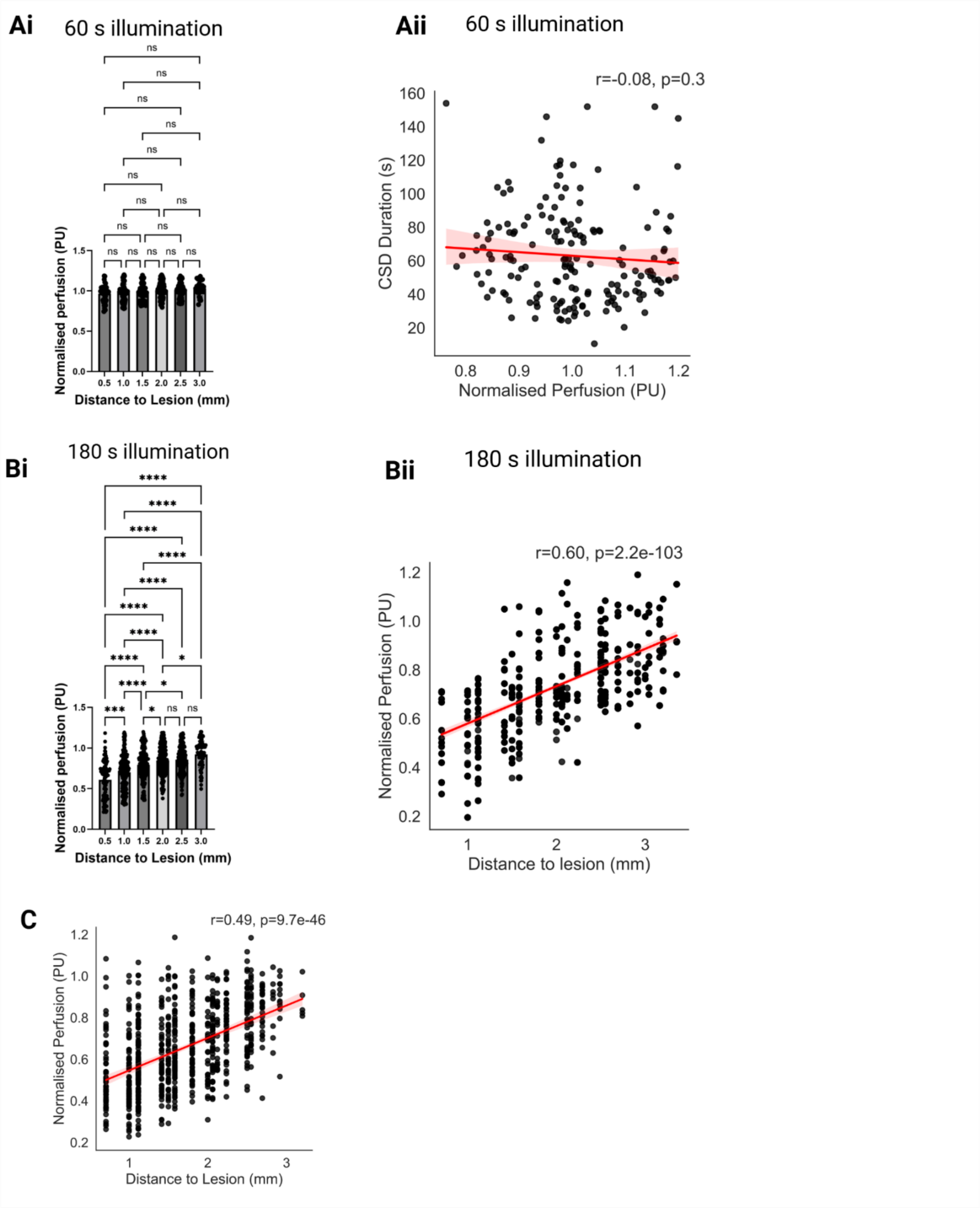


Supplementary figure 4. **Ai**. Normalised perfusion (PU) as a function of distance to the lesion (mm) after 60 second photothrombosis (0.97±0.11 PU at illuminated region). **Aii**. Quantified CSD duration (s) compared against normalised perfusion (PU) post initial 60 second insult (Spearman's, r = -0.08, p = 0.3, n =15). **Bi.** Grouped analysis of mean CSD duration (s) as a function of normalised perfusion (PU), revealing significant differences across most perfusion bins (mean ± s.d.). **Bii**. Plots of normalised perfusion (PU) as a function of distance to the lesion (mm) after 180 second photothrombosis (Spearman's, r = 0.60, p = 2.2e-103, n =15). **C**. Plot of normalised perfusion (PU) with distance to the ischaemic lesion core (mm) from dMCAo mice 1 hr post ischaemia induction (Spearman's, r = 0.49, p = 9.7e-46, n =15).

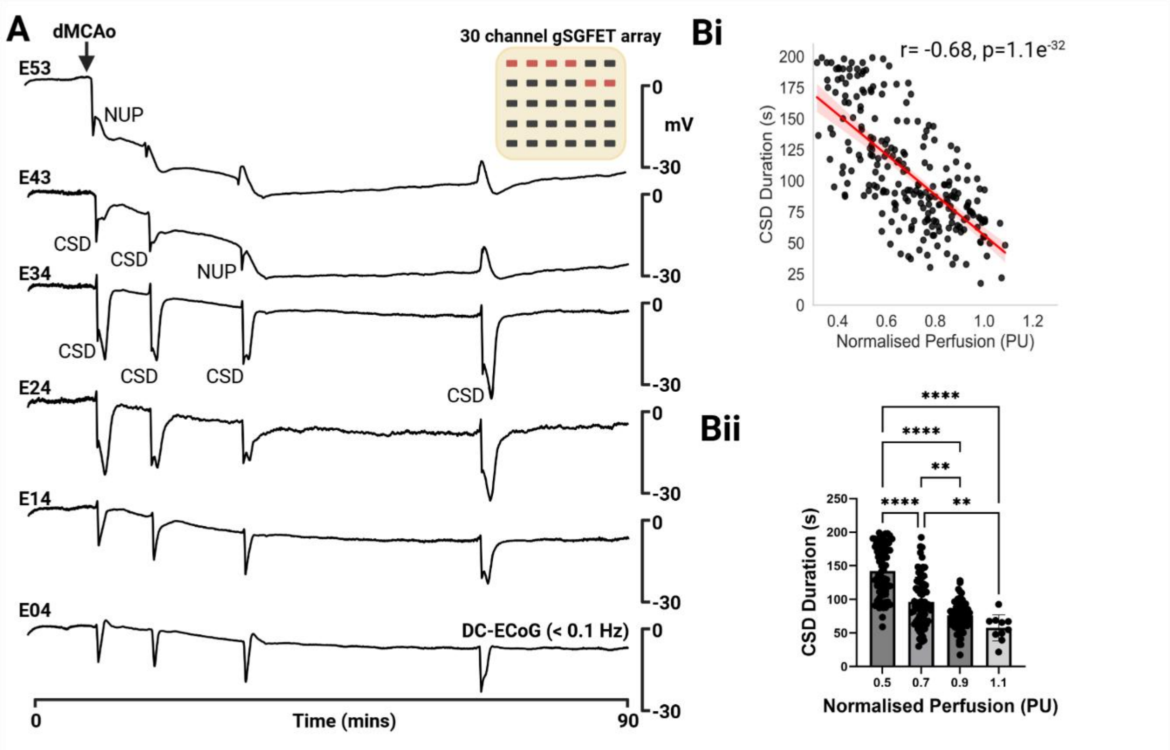


Supplementary figure 5. **A**. Example 90 min DC-coupled (DC-ECoG < 0.1 Hz) gSGFET recording following dMCAo in anaesthetised mice. Channels have been selected at increasing distances from the lesion core, represented in the array map. Note the CSD duration changing with distance. **Bi**. Scatter plot displaying a negative correlation between CSD duration (s) and normalised perfusion (PU) in the dMCAo model ($r = -0.68$, $p=1.1e^{-32}$). **Bii**. Bar graph showing average CSD duration (s) as a function of binned normalised perfusion (PU). Group-wise comparison of normalised perfusion against CSD duration allowed for further statistical comparison and displayed significant differences across most perfusion groups 1 (One-way ANOVA, **** $p < 0.0001$, ** $p < 0.01$, n = 15), except for 0.9 vs 1.1. Mean CSD durations were 142.2±38.6 s at 0.5 PU, 96.3±36.7 s at 0.7 PU, 75.8±22.1 s at 0.9 PU and 57.6±19.4 s at 1.1 PU.

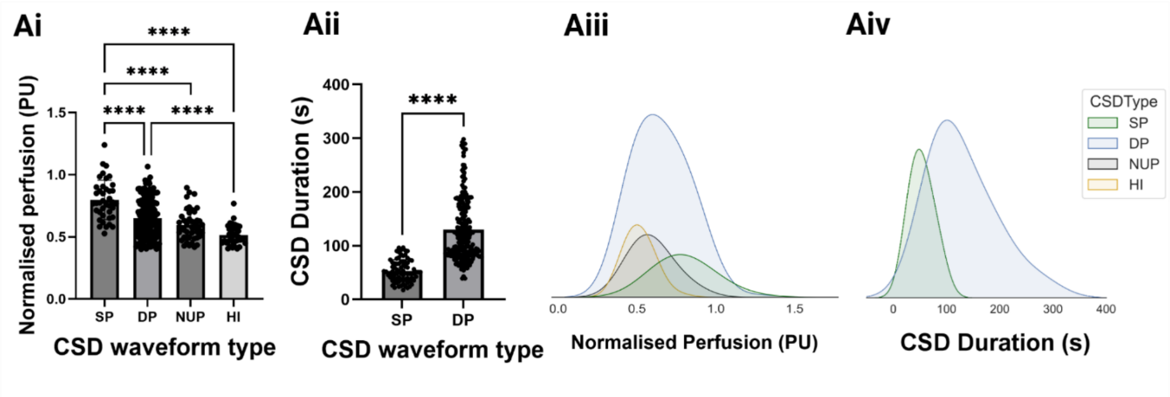


Supplementary figure 6. **Ai**. Plot comparing CSD waveform types (40 SP, 170 DP, 44 NUP, and 35 HI) recorded following dMCAo in mice against normalised perfusion (PU) (One-way ANOVA, **** $p < 0.0001$, n = 15). **Aii**. Plot showing DP (223) and SP (78) CSD duration (s) (unpaired t-test, **** $p < 0.0001$, n = 15). **Aiii**. Density plot displaying CSD type distribution with normalised perfusion (PU). **Aiv**. Density plot displaying DP and SP CSD duration (s) distribution.

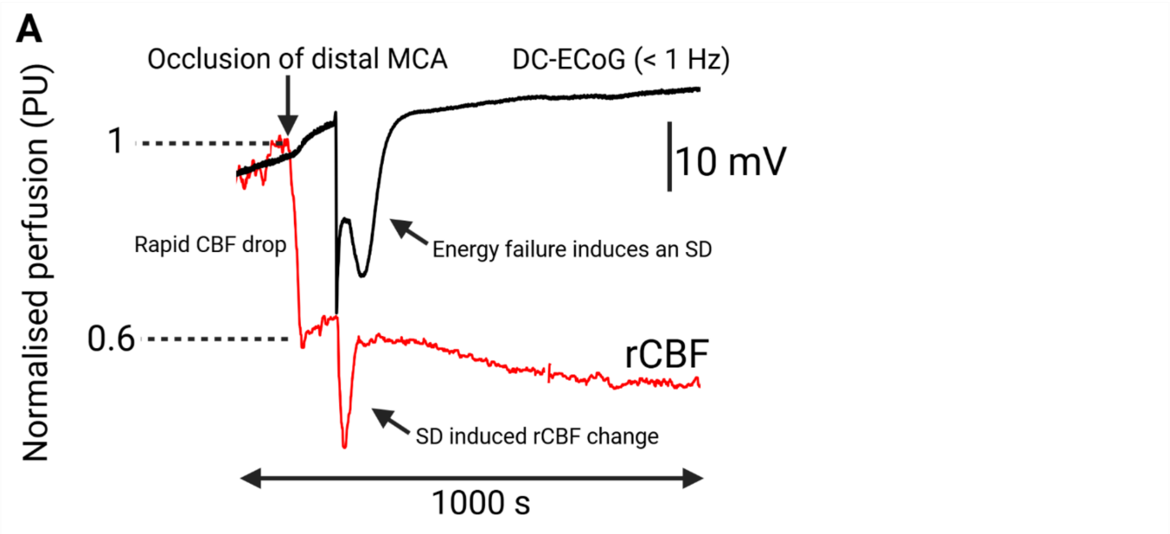


Supplementary figure 7. Schematic displaying the order of ischaemia-induced CSD and CSD induced haemodynamic response. A rapid CBF drop causes energy failure and induces a CSD. The CSD then induces a haemodynamic response.

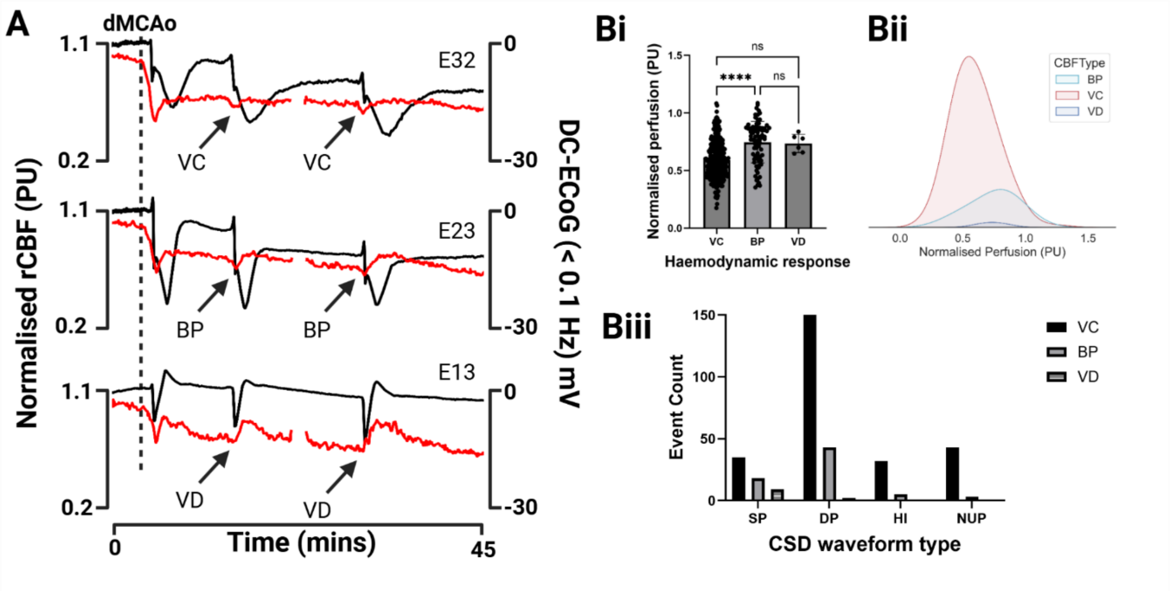


Supplementary Figure 8. Cortical spreading depolarisation induced haemodynamic responses are dependent on regional cerebral blood flow **A**. Example DC-ECoG and rCBF traces post dMCAo in anaesthetised mice. Selected channels are at increasing distance (0.5 mm apart) from the lesion core. Three haemodynamic responses were observed in this work, vasoconstriction (VC), biphasic response (BP) and vasodilation **Bi-ii**. CSD-induced haemodynamic response type plotted in function of mean normalised perfusion (PU) (298 VC responses at 0.59±0.17 PU, 78 BP responses at 0.75±0.18 PU and 6 VD responses at 0.74±0.08 PU) and presented as a density distribution plot from dMCAo induced mice. **Ciii**. Count of CSD-induced haemodynamic responses with relation to CSD waveform type (SP induced 35 VC, 18 BP and 9 VD; DP induced 150 VC, 43 BP and 2 VC; HI induced 32 VC, 5 BP and NUP induced 43 VC and 3 BP).

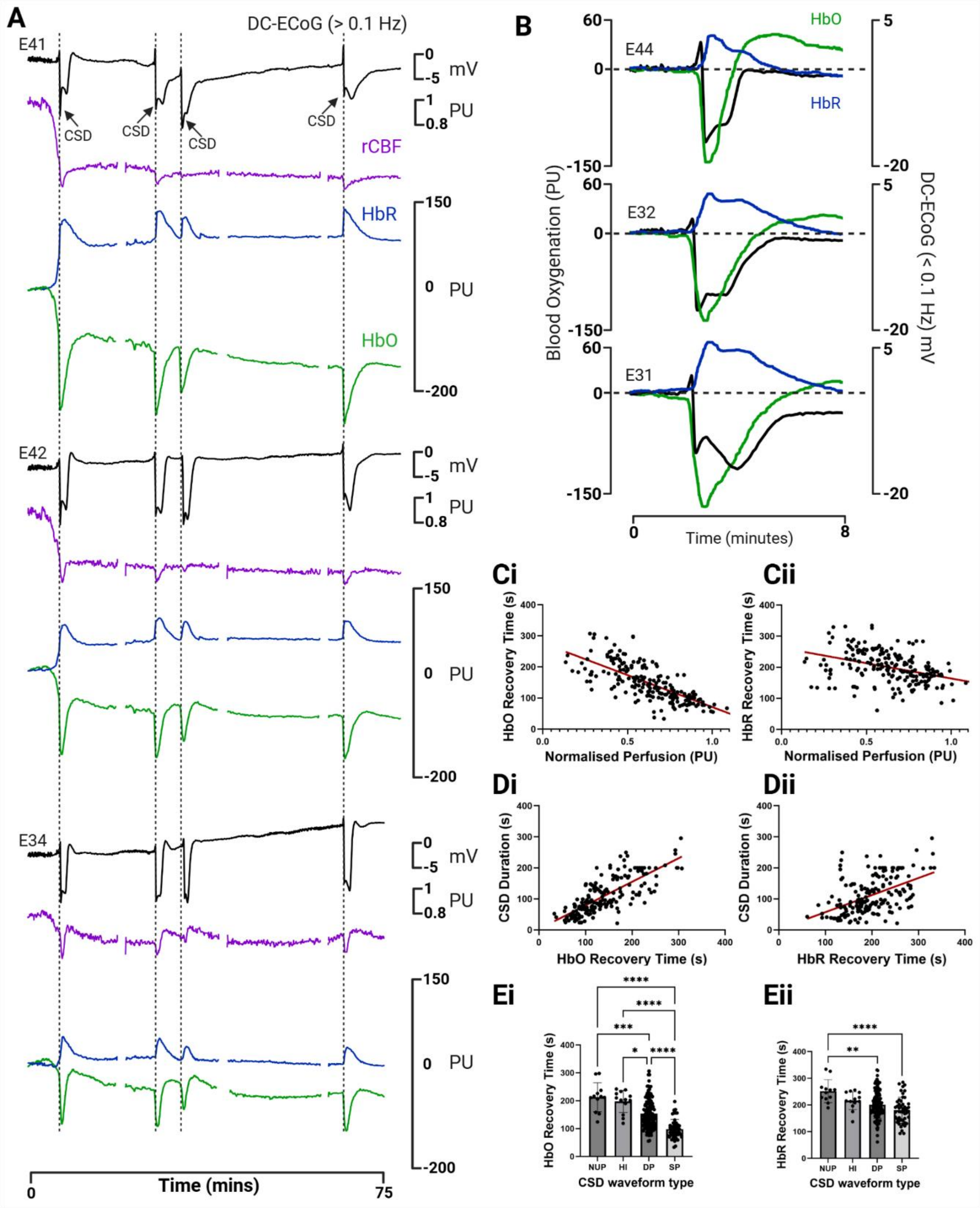


Supplementary Figure 9. CSD duration and waveform are predictive of transient tissue oxygenation deficits. **A**. Example 75-minute DC-ECoG recording (> 0.1 Hz), using a gSGFET array, with concurrent regional cerebral blood flow (rCBF (PU)) and blood oxygenation (HbO/ HbR (PU)) imaging. Dotted lines marks the start of CSDs. **B**. Example zoomed in DC-ECoG recording (8 minutes) with simultaneous blood oxygenation (HbO/ HbR (PU)) imaging. **Ci-Cii**. Quantification of HbO and HbR recovery time (s) plotted against normalised regional cerebral blood flow (PU) (HbO linear regression, R squared = 0.54, $p < 0.0001$; HbR linear regression, R squared = 0.16, $p < 0.0001$) . **Di-Dii**. Plot displaying CSD duration (s) in function of HbO and HbR recovery time (s). Bar plot displaying HbO and HbR recovery time (s) and CSD waveform type.
